# The transcriptional cofactor IRF2BP2 plays a key role in T cell homeostasis and Treg cell expansion

**DOI:** 10.1101/2021.07.29.454283

**Authors:** Giuliana P. Mognol, Barbara Oliveira-Vieira, Natalia Pinheiro-Rosa, Barbara C. Peixoto, Marianna Boroni, Edahi González-Avalos, Cristiane Secca, Hozefa Bandukwala, Ana Maria C. Faria, Anjana Rao, João P.B. Viola

**Affiliations:** Program of Immunology and Tumor Biology, Brazilian National Cancer Institute (INCA), Rio de Janeiro, RJ, Brazil; Division of Signaling and Gene Expression, La Jolla Institute for Immunology, La Jolla, CA 92137; Department of Biochemistry and Immunology, Institute of Biological Science, Federal University of Minas Gerais, Belo Horizonte, MG, Brazil; Laboratory of Bioinformatics and Computational Biology, Brazilian National Cancer Institute (INCA), Rio de Janeiro, RJ, Brazil; Bluestar Genomics, La Jolla, CA, USA; Department of Pathology and Immunology, Washington University School of Medicine, St. Louis, USA; Sigilon Therapeutics, Cambridge, MA, USA

**Author notes:** **Corresponding author:** Programa de Imunologia e Biologia Tumoral; Instituto Nacional de Câncer (INCA); Rua André Cavalcanti, 37; Centro, Rio de Janeiro, RJ 20231-050, Brazil. The first two authors contributed equal in this work.

**Keywords:** IRF2BP2, IFN-γ, FoxP3, Treg

## Abstract

The levels of the co-transcriptional regulator IRF2BP2 (Interferon Regulatory Factor-2 Binding Protein-2) decrease with T cell activation and, when ectopically expressed, it reduces T cell proliferation. To further characterize the function of IRF2BP2 in T cell responses in vivo, we generated a conditional transgenic knock-in mouse that overexpresses IRF2BP2 in T lymphocytes. Overexpression of IRF2BP2 leads to a reduction in the T cell compartment of naive animals, upregulation of *Foxp3* and *Ifng*; an increase in the frequency of regulatory T cells (Tregs), a preferential Th1 differentiation with increase of IFN-γ production and a reduction of T cell proliferation, suggesting a disruption in T cell homeostasis. Interestingly, knock-in mice displayed reduced clinical and inflammatory signs of Experimental Autoimmune Encephalomyelitis (EAE) when compared to the control mice, with an augmented frequency of Treg cells. Altogether, our findings indicate that IRF2BP2 might help to control exacerbated T cell responses and point to a role for IRF2BP2 in preventing T cell autoimmunity.

## Introduction

The IRF (Interferon Regulatory Factor) family of transcription factors are regulators of type I interferons in response to viral infection (Lohoff *et al*., 2000), important for T helper cell differentiation and essential in regulating immune response, cell survival and oncogenesis (Tamura *et al*., 2008; Savitsky *et al*., 2010). IRF-2 downregulates type I IFN-induced genes by antagonistically repressing IRF-1-binding activity (Savitsky *et al*., 2010). Identified in 2003 as a nuclear transcriptional corepressor of IRF-2, IRF2BP2 (Interferon Regulatory Factor-2 Binding Protein-2) (Childs & Goodbourn, 2003) was further described as an anti-apoptotic (Koeppel *et al*., 2009; Yeung *et al*., 2011) and pro-survival factor (Koeppel *et al*., 2009; Tinnikov *et al*., 2009), also involved in cellular differentiation (Stadhouders *et al*., 2015). The majority of studies depict IRF2BP2 as a co-transcriptional repressor (Childs & Goodbourn, 2003; Koeppel *et al*., 2009; Tinnikov *et al*., 2009, Carneiro *et al*., 2011, Yeung *et al*., 2011; Stadhouders *et al*., 2015), but there are descriptions of IRF2BP2 as a transcriptional activator, activating VEGF-A and favoring revascularization after ischemia (Teng *et al*., 2010, Chen *et al*., 2015). For a detailed review of IRF2BP2 structure and functions, please refer to Ramalho-Oliveira, 2019 (Ramalho-Oliveira *et al*., 2019).

IRF2BP2 has been implicated in immune responses, interacting and repressing the functions of the transcription factor NFAT1 (Carneiro *et al*., 2011); repressing/delaying lymphocyte activation (Carneiro *et al*., 2011, Secca *et al*., 2016) and proliferation (Secca *et al*., 2016); and modulating macrophage polarization towards the anti-inflammatory, M2 phenotype (Chen *et al*., 2015). Translocations and mutations in the human *irf2bp2* gene have been observed in various cancers including primary central nervous system lymphoma (Pall & Hamilton, 2008), multiple myeloma (Ni *et al*., 2012), mesenchymal chondrosarcoma (Nyquist *et al*., 2012) and acute promyelocytic leukemia (Yin *et al*., 2015), emphasizing the importance of the tight control of IRF2BP2 expression, which are yet poorly understood as are the pathways regulated by IRF2BP2 in normal cell physiology.

IRF2BP2 expression is reduced in T cells of patients with Multiple Sclerosis and restored to normal levels in patients in clinical remission, suggesting a role for IRF2BP2 in T cell-mediated inflammation (Arruda *et al*., 2015). Multiple Sclerosis results from failure of tolerance mechanisms such as T regulatory (Treg) cells to prevent the expansion of pathogenic T cells. Treg cells express FOXP3, a transcription factor required for their development and suppressive function (Fontenot *et al*., 2005, Zheng & Rudensky, 2007) and are required for immune self-tolerance. Therefore, dysfunctional Tregs (Viglietta *et al*., 2004) and abnormalities in FOXP3 expression (Huan *et al*., 2005) are important in the pathogenesis of Multiple Sclerosis (Arruda *et al*., 2015). In mice, Experimental Autoimmune Encephalomyelitis (EAE) is a model of autoimmune disease that mimics what occurs in patients with multiple sclerosis in which CD4^+^ T cells are critical players in the inflammatory process affecting the central nervous system (Miller *et al*., 2010; Rangacharia & Kuchroo, 2013). IFN-γ is an important cytokine during the progression of Multiple Sclerosis and EAE (Arellano *et al*.,1995) and it has been discussed that changes in the regulatory factors that affect the feedback control of multiple IFN-stimulated genes lead to aberrant cytokine response and inflammation (Arruda *et al*., 2015).

To better understand the effects of IRF2BP2 during T cell activation and differentiation and to evaluate its functions in vivo, we constructed and characterized a conditional IRF2BP2 transgenic mice that overexpress IRF2BP2 in T lymphocytes (IRF2BP2^fl/fl^Lck-cre^+^). These mice developed normally but exhibited an increased Treg (CD4^+^CD25^+^Foxp3^+^) population, and a higher expression of FOXP3 and IFN-γ when compared to control mice. Transgenic IRF2BP2^fl/fl^Lck-cre^+^ CD4^+^ T cells, when differentiated *in vitro* to either Th1, Th2 or Th17, produced more IFN-γ than control cells, underscoring the role of IRF2BP2 as an important regulator of IFN-γ expression. Making use of an induced EAE model, we observed that the clinical signals of the disease were milder in mice overexpressing IRF2BP2 versus control mice, with less inflammatory T CD4^+^ cell infiltrate in the spinal cord. Splenic cells from IRF2BP2 knock-in mice also exhibited an increased number of Tregs, suggesting that the reduction in the symptoms of EAE disease was likely due to an increase in immunoregulatory activity. Taken together, our results demonstrate a function for IRF2BP2 in T cell effector response and further suggest a regulatory role for IRF2BP2 in the control of autoimmunity.

## Results

### Construction and characterization of an IRF2BP2 conditional knock-in mice

We constructed a conditional transgenic mouse that overexpressed IRF2BP2 by inserting the coding region of the *Irf2bp2* gene into the ROSA26 locus, a locus that codifies an ubiquitously expressed non-coding RNA (see *Methods*). A schematic representation of the transgenic cassette inserted into the ROSA26 locus is shown in **Figure 1A**. It contains a neomycin resistant gene (NeoR) and a stop codon flanked by loxP sites, making the expression of transgenic IRF2BP2 conditional to the presence of Cre recombinase. The transgene also contains an internal ribosomal entry site (IRES) followed by the EGFP sequence, which is expressed alongside with IRF2BP2 and works as a reporter gene. The CTV vector containing the transgene cassette was linearized and electroporated into embryonic stem (ES) cells. After 3 weeks, gDNA of the G418-resistant ES colonies was extracted, and the transgene integration was confirmed by both PCR (*data not shown*) and Southern blot (**Figure 1B**). Based on the EcoRI digestion pattern shown in **Figure 1A**, the wild type allele has 11.6 Kb and the transgene allele has 5.9 Kb. Two clones containing the transgene allele were injected into albino C57BL/6 blastocysts to generate chimerical mice, which were then bred with C57BL/6 mice for germline transmission. The IRF2BP2^fl/fl^ mice generated were bred with Lck-cre^+^ mice to generate the IRF2BP2^fl/fl^Lck-cre^+^ mice, which overexpresses IRF2BP2 in T lymphocytes, as shown in **Figure 1C**.

**Figure 1.**
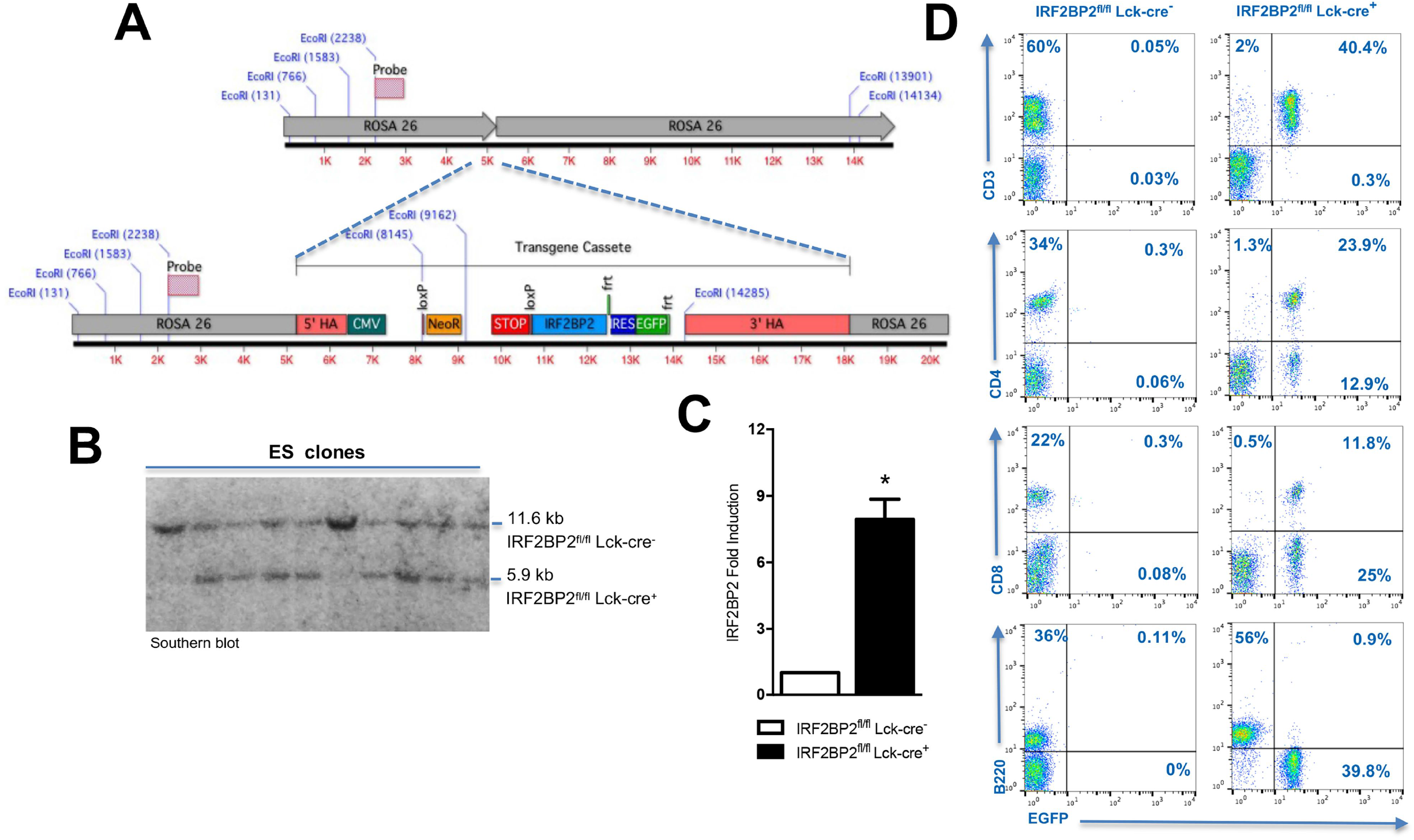
Construction and expression of a conditional knock-in IRF2BP2 mice. **A**. Schematic representation of the transgene inserted into the ROSA26 locus to produce the conditional IRF2BP2 knock-in mice. Top: Wild type ROSA26 locus showing the EcoRI digestion pattern. Bottom: ROSA26 locus with the transgene inserted, showing the EcoRI digestion pattern. The place where the Southern blot probe anneals is indicated. **B**. Southern blot result with different G418-resistant embryonic stem cell clones obtained after 3 weeks of electroporation with the CTV-mIRF2BP2-myc plasmid. Based on the EcoRI digestion pattern showed in **A**, the wild type allele has 11.6 Kb and the transgene allele has 5.9 Kb. Two clones with the transgene allele were injected into albino C57BL/6 blastocysts to generate chimerical mice, which were bred with C57BL/6 mice for germline transmission. The IRF2BP2^fl/fl^ mice generated were bred with Lck-cre^+^ mice to generate the IRF2BP2^fl/fl^Lck-cre^+^ mice. **C**. Real Time PCR showing the relative quantification of *Irf2bp2* mRNA expression in CD4^+^ T cells from IRF2BP2^fl/fl^Lck-cre^+^ mice compared to IRF2BP2^fl/fl^Lck-cre^-^ mice. The cells were stimulated for 48 hours with anti-CD3 and anti-CD28. The *Irf2bp2* gene expression was normalized by the HPRT expression. Representative of 4 independent experiments (n=4). **D**. Flow cytometry showing GFP expression in B^+^, CD4^+^ or CD8^+^ cells obtained from lymph nodes of IRF2BP2^fl/fl^Lck-cre^+^ or from IRF2BP2^fl/fl^Lck-cre^-^ mice. Representative of 2 independent experiments (n=3).

Using GFP as a reporter to evaluate the overexpression of IRF2BP2 in lymphocytes from lymph nodes of IRF2BP2^fl/fl^Lck-cre^+^ and IRF2BP2^fl/fl^Lck-cre^-^ mice, we observed, as expected, that T cells (either CD3^+^, or CD4^+^ and CD8^+^) but not B cells from the IRF2BP2^fl/fl^Lck-cre^+^ mice, express GFP (**Figure 1D**). IRF2BP2 is described as a nuclear protein (Childs & Goodbourn, 2003; Carneiro *et al*., 2011), and confocal microscopy analysis of transgenic CD4^+^ T cells activated for 24 hours with PMA and ionomycin shows that the IRF2BP2 protein expressed by the transgene behaved as the endogenous protein in terms of cellular localization (data not shown).

### Mice overexpressing IRF2BP2 exhibit a reduced frequency of T cells in the peripheral lymphoid organs, and a higher percentage of Tregs

As the IRF2BP2^fl/fl^Lck-cre^+^ mice overexpress IRF2BP2 in T cells, we were interested in evaluate the T cell thymic differentiation (**Figure 2A-B**). In fact, there was no difference in the frequency of double negative, double positive, single CD4^+^ or single CD8^+^ cell populations between IRF2BP2^fl/fl^Lck-cre^+^ and IRF2BP2^fl/fl^Lck-cre^-^ mice, indicating normal thymic differentiation. We then evaluated lymph nodes and splenic T and B cell populations of these mice. In all our experiments, the percentage of T cells (either CD3^+^, CD4^+^ or CD8^+^) was reduced, while the percentage of B cells was increased in lymphocytes isolated from IRF2BP2^fl/fl^Lck-cre^+^ mice compared to the control mice (**Figure 2C-F** and **Figure 1D**), in accordance with our previous report showing that IRF2BP2 is a repressor of proliferation (Secca *et al*., 2016 and unpublished data). Mice overexpressing IRF2BP2 had a significant increase in the frequency of Tregs (CD4^+^CD25^+^FoxP3^+^) when compared to control mice, both in the thymus and in the periphery (**Figure 3**), what could at least in part be responsible for the smaller number of T cells found on those mice.

**Figure 2.**
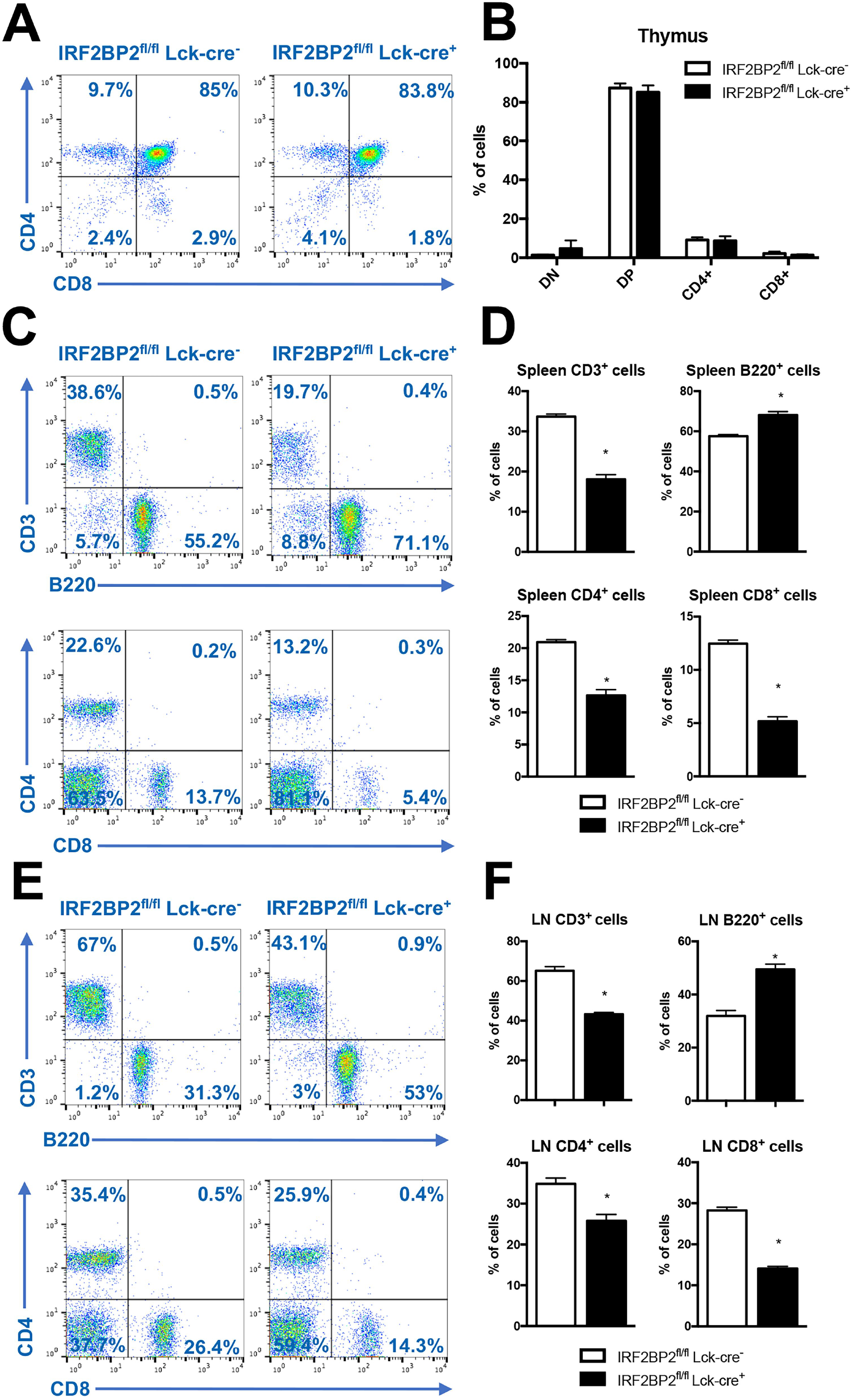
Percentage of B and T cell populations of IRF2BP2^fl/fl^Lck-cre^+^ and IRF2BP2^fl/fl^Lck-cre^-^ mice. The thymus (**A-B**), spleen (**C-D**) and lymph nodes (**E-F**) were taken from the indicated mice. **A, C, E**. Flow cytometry of the indicated organs stained for anti-B220, - CD3, -CD4 and -CD8, as indicated. **B, D, F**. Percentage of the indicated cell population relative to the graph shown on the left. The bar graphs show average ± SD (n=6). (*) p<0.05. LN-Lymph nodes; DN – double negative cells; DP – double positive cells. Representative of 2 independent experiments (n=6).

**Figure 3.**
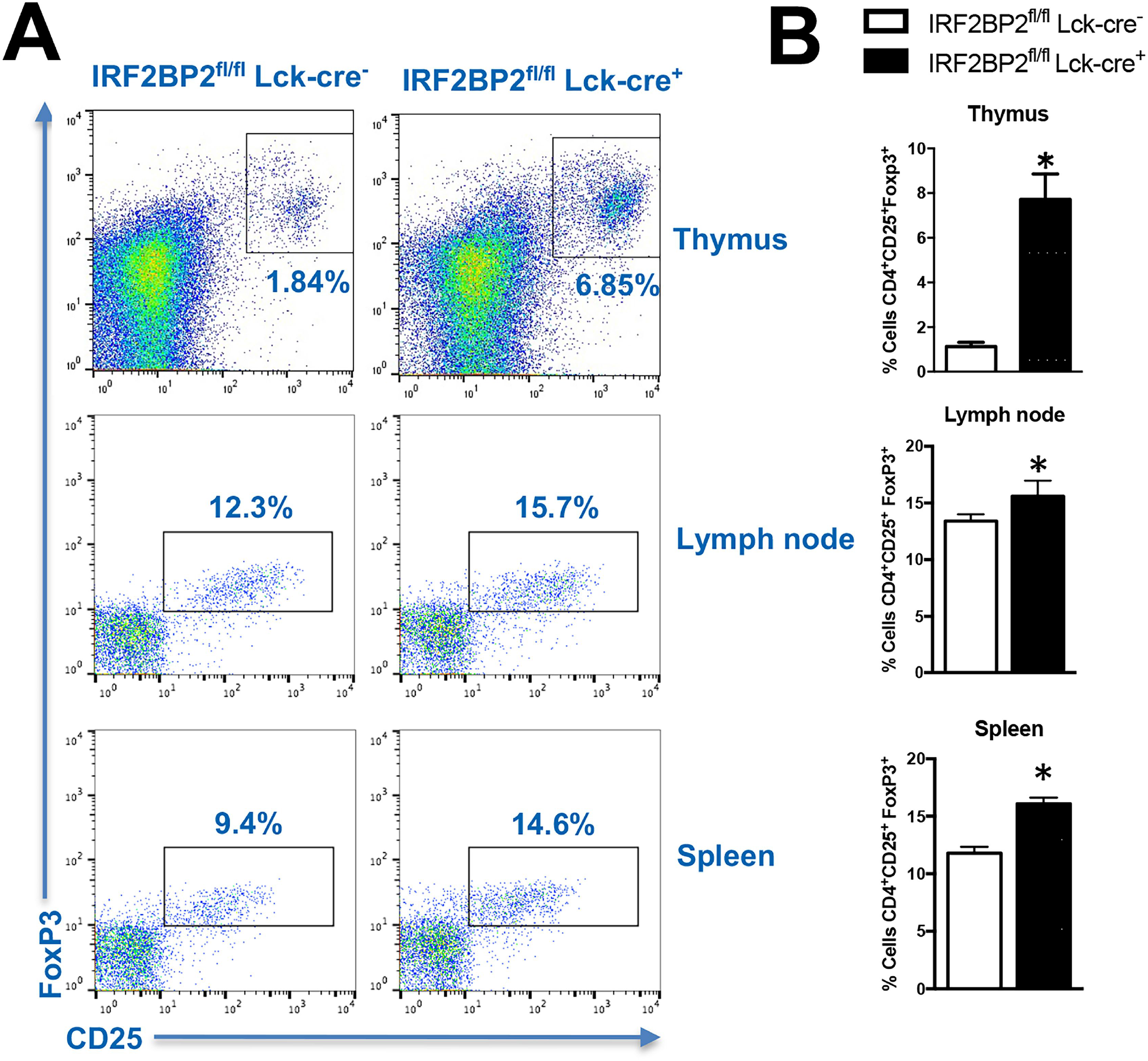
Percentage of Treg (CD4^+^CD25^+^FoxP3^+^) cells in central and peripheral immune organs of IRF2BP2^fl/fl^Lck-cre^+^ and IRF2BP2^fl/fl^Lck-cre^-^ mice. Lymphocytes from the thymus, spleen or lymph nodes of the indicated mice were stained for anti-CD4, anti-CD25 and anti-FoxP3. **A**. Flow cytometry displaying the percentage of Tregs. **B**. Bar graphs showing the average of Tregs ± SD relative to the graph shown in (**A**) (n=6). (*) p<0.05. Representative of 2 independent experiments (n=6).

### Expression of *Ifng* and *Foxp3* is upregulated in IRF2BP2^fl/fl^Lck-cre^+^ CD4^+^ T cells

IRF2BP2 has been described as a co-transcriptional regulator (Childs & Goodbourn, 2003; Koeppel *et al*., 2009, Tinnikov *et al*., 2009, Carneiro *et al*., 2011, Yeung *et al*., 2011; Stadhouders *et al*., 2015). To elucidate the transcription profile regulated by IRF2BP2, we performed an RNA-seq experiment with sorted naïve CD4^+^ T cells from IRF2BP2^fl/fl^Lck-cre^+^ and IRF2BP2^fl/fl^Lck-cre^-^ mice, which were stimulated for 1 hour with PMA plus ionomycin or left unstimulated (see *Methods*) (**Figure 4**). 159 genes were upregulated and 45 genes were downregulated in CD4^+^ T cells from IRF2BP2^fl/fl^Lck-cre^+^ mice (**Figure 4A** and **Dataset 1**). A genome browser view showing IRF2BP2 expression in unstimulated (US) and PMA/ionomycin-stimulated cells is shown in **Figure 4B**. The differentially expressed genes important to T cell biology fell into several categories, including cytokines (*Ccl4, Ccl5, Ifng, Xcl1*), transcription factors (*Foxp3, Irf2bp2, Pou2af1, Tcf4*), cell surface receptors involved in both T cell activation (*Tnfrsf9 [4-1BB], Tnfrsf4 [OX40])* and inhibition (*Pdcd1 [PD-1]*), and chemokine receptors (*Cxcr3, Cxcr5, Cxcr6*). The downregulated genes include *Fosb*, a member of AP-1 family of transcription factors, implicated in the regulation of T cell activation and proliferation (Shaulian & Karin, 2001) and proteins of the Krüppel-like factor family of transcription factors (*Klf4, Klf*6 and *Klf*10), zinc finger transcription factors involved in the differentiation, activation, quiescence, or homing of various T-cell subsets (Cao *et al*., 2010) (**Figure 4C**). Among these genes, we focused on the upregulation of *Ifng* and *Foxp3*, whose relevance will be discussed in the upcoming sections.

**Figure 4.**
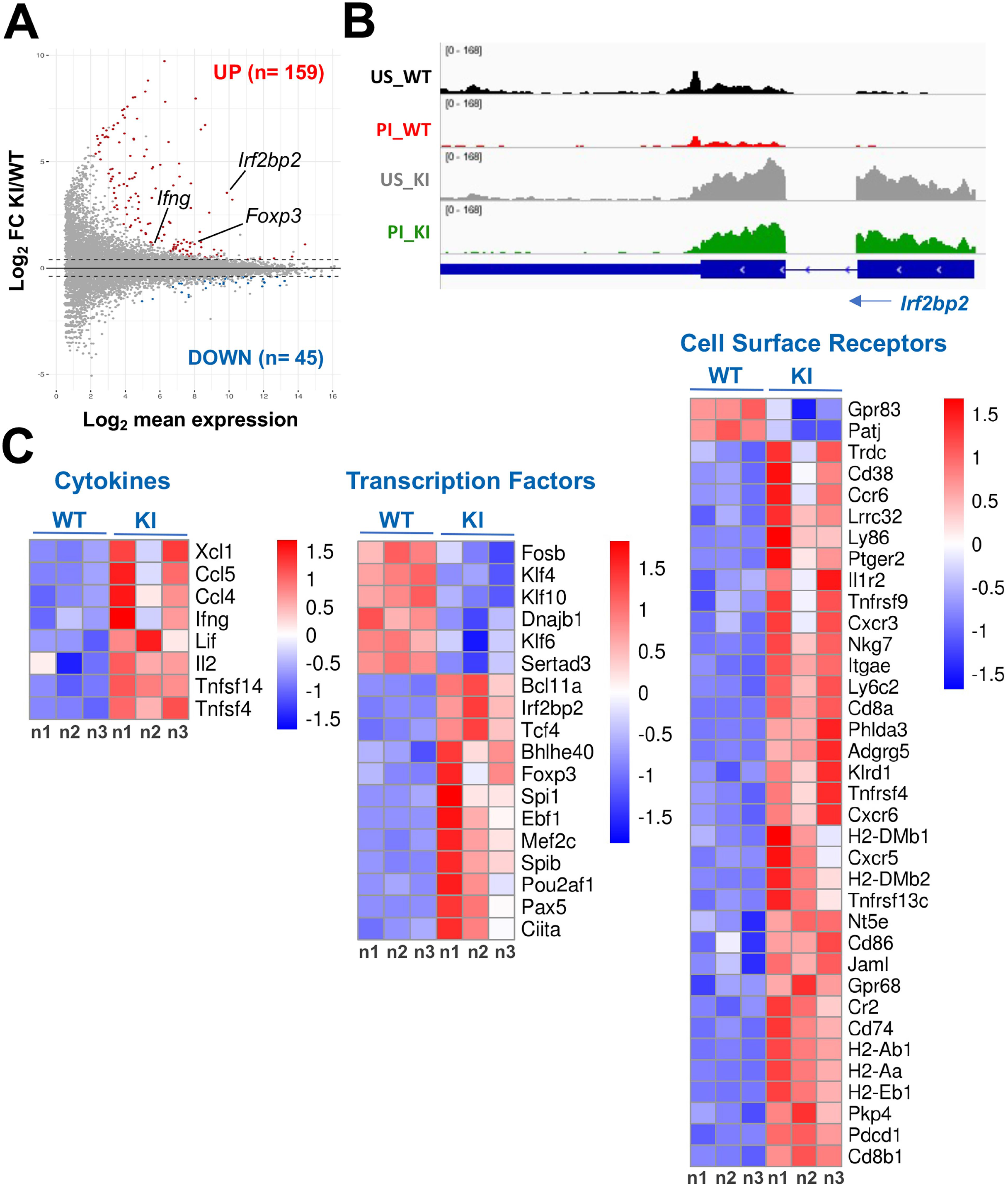
Transcriptional program regulated by IRF2BP2 in CD4^+^ T lymphocytes. The RNA-seq was performed with sorted naïve CD4^+^ T cells from IRF2BP2^fl/fl^Lck-cre^+^and IRF2BP2^fl/fl^Lck-cre^-^ mice. **A**. MA plot showing the log2-fold change of mRNA transcript levels and mean expression between IRF2BP2^fl/fl^Lck-cre^+^and IRF2BP2^fl/fl^Lck-cre^-^ control cells, *in vitro* stimulated for 1 hour with PMA (10 nM) and ionomycin (1 μM). Each dot represents an expressed gene. The number or genes significantly upregulated or downregulated in IRF2BP2^fl/fl^Lck-cre^+^ versus IRF2BP2^fl/fl^Lck-cre^-^ cells is indicated (log2FC ≥ 0.4 and padj ≤ 0.05). **B**. Genome browser view of RNA-seq signal at the *Irf2bp2* locus in the different experimental conditions. Blue boxes: exons. The arrow indicates the transcription direction. **C**. Heatmaps representation of log2-fold changes in gene expression in IRF2BP2^fl/fl^Lck-cre^+^ compared to IRF2BP2^fl/fl^Lck-cre^-^ cells, showing individual replicates. US – unstimulated cells, PI – PMA + ionomycin stimulated cells, WT – IRF2BP2^fl/fl^Lck-cre^-^ cells, KI – IRF2BP2^fl/fl^Lck-cre^+^cells.

### A substantial number of *in vitro* differentiated IRF2BP2^fl/fl^Lck-cre^+^ CD4^+^ T cells produce IFN-γ, regardless of their effector profile

To evaluate the effect of IRF2BP2 on CD4^+^ T cell differentiation, we *in vitro* activated and skewed CD4^+^ T cells towards Th1, Th2 and Th17 effector profiles (**Figure 5A**). On day 6, we re-stimulated the cells with PMA and ionomycin to quantify the production of the main cytokines related to each profile. On one hand, wild type cells, as expected, produced IFN-γ, but not IL-4 when differentiated to Th1; IL-4 but not IFN-γ when differentiated to Th2 and IL-17 but not IFN-γ when differentiated to Th17 (**Figure 5B-C, left panels**). On the other hand, a small percentage of cells overexpressing IRF2BP2 produced IFN-γ regardless of the differentiated phenotype (**Figure 5B-C, right panels**). When differentiated to Th1, a higher percentage of cells overexpressing IRF2BP2 produced IFN-γ compared to control cells (78.7% versus 69.3%, respectively, **Figure 5B-C**). Remarkably, even though anti-IFN-γ is present in the cultures undergoing differentiation to Th2 cells, 18.7% of the IRF2BP2^fl/fl^Lck-cre^+^ cells differentiated under Th2-polarising conditions produced IFN-γ, with 11.4% of those producing both IL-4 and IFN-γ. The same was observed with cultures differentiated to Th17, which also contain anti-IFN-γ: in this case, 14.2% of the IRF2BP2^fl/fl^Lck-cre^+^ cells produced IFN-γ, 5.7% of those producing both IL-17 and IFN-γ (**Figure 5B-C**).

**Figure 5.**
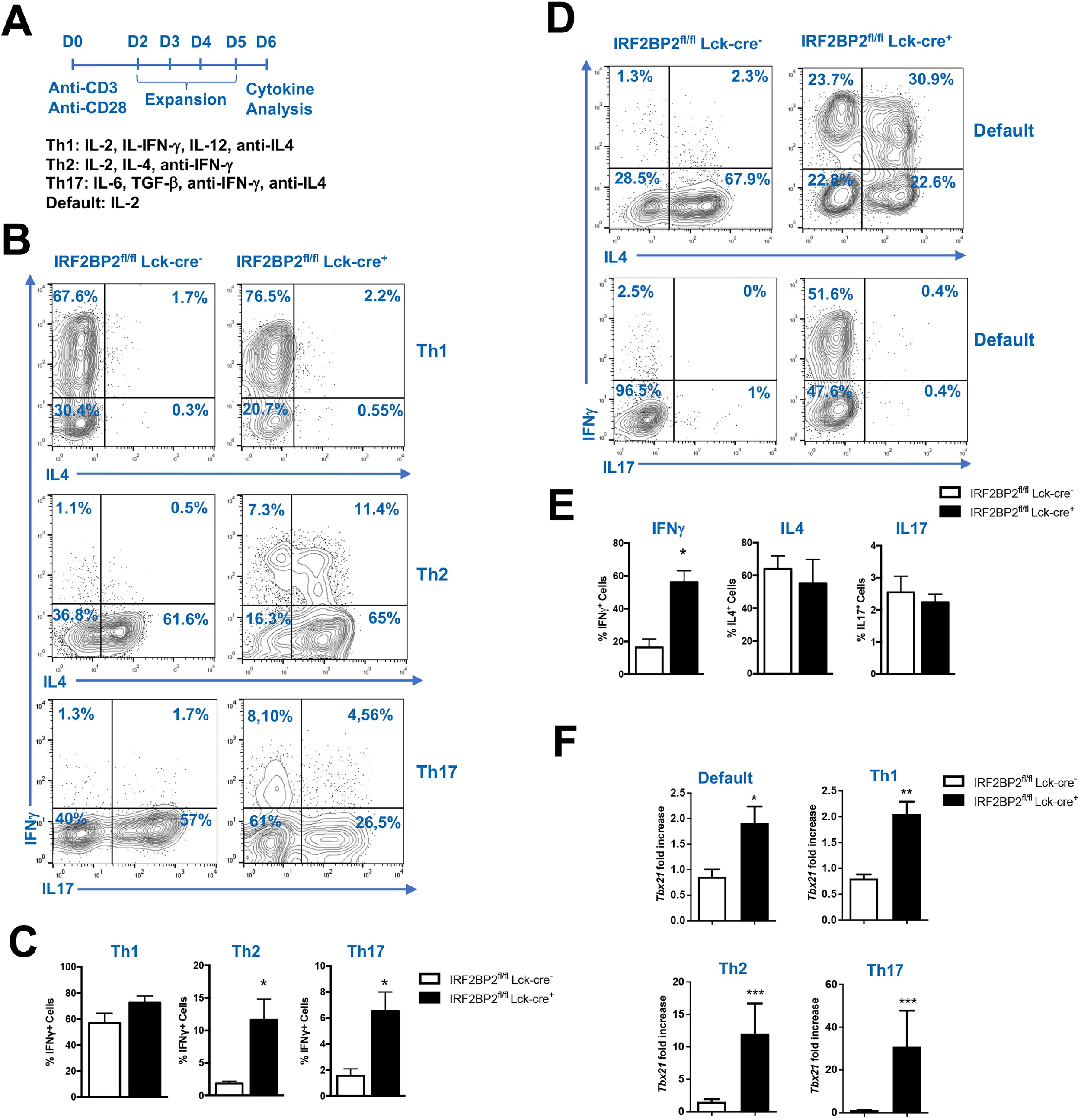
*In vitro* CD4^+^ T cell differentiation to the effector profiles. **A**. Schematic representation of the experiment. CD4^+^ T cells purified from lymph nodes of IRF2BP2^fl/fl^Lck-cre^+^ or IRF2BP2^fl/fl^Lck-cre^-^ mice were stimulated with anti-CD3 (1 μg/mL) and anti-CD28 (1 μg/mL) in the presence of the indicated cytokines and were differentiated to Th1, Th2, Th17 or default, as indicated. After 6 days, the cells were stimulated for 6 hours with PMA (10 nM) and ionomycin (1 μM) and stained for anti-IFN-γ, -IL-4 and -IL-17, depending on the differentiation profile. **B, D**. Flow cytometry showing the intracellular cytokine staining with the specific antibodies indicated for each profile. **C**. The dot plots show the percentage of IFN-γ producing cells in each profile. **E**. Percentage of cells producing IFN-γ, IL-4 and IL-17 in the default differentiation. Graphs show average ± SD. (*) p<0.05. Representative of 3 independent experiments (n=5). **F**. T-bet *(Tbx21)* mRNA fold induction was quantified by RT-PCR on purified CD4^+^ T cells differentiated to Th1, Th2, Th17 and default, as indicated. The data refer to purified cells compared to the lower mean obtained for IRF2BP2^fl/fl^Lck-cre^-^ *Tbx21* expression. For relative quantification the mass of cDNA used for the reaction was normalized using the *Hprt* housekeeping gene. Bar graphs showing the average ± SD. The (*) p<0.05, and (***) p<0.0001. Representative of 4 independent experiments (n=11).

We then cultured CD4^+^ T cells stimulated with anti-CD3 and anti-CD28 in the absence of polarizing cytokines (default differentiation) and re-stimulated them on day 6 with PMA and ionomycin to access cytokine production (**Figure 5D-E**). Wild type cells mainly produced IL-4 (∼ 70%), without any production of IFN-γ (**Figure 5D-E**), indicating that these cells differentiate to Th2 phenotype by default, whereas ∼50% of the IRF2BP2^fl/fl^Lck-cre^+^ cells produced IL-4 and ∼50% produced IFN-γ, 30% of those producing both IL-4 and IFN-γ (**Figure 5D-E**). This IL-4^+^IFN-γ^+^ double positive population displayed a Th0 profile, defined as cells that produce both Th1 and Th2 cytokines (Yates, 2004), normally found at the beginning of T cell differentiation in recently activated cells (Murphy *et al*., 2002), suggesting an incomplete or a delayed Th differentiation.

All these findings suggest that the mechanisms of IFN-γ repression are deregulated in IRF2BP2^fl/fl^Lck-cre^+^ CD4^+^ T cells. To further explore the reason for the higher production of IFN-γ by the transgenic T cells in comparison to control cells, we evaluated the expression of transcription factors important for Th cell differentiation and found that T-bet, a transcription factor known to bind to, increase histone acetylation and transactivate the IFN-γ promoter (Szabo *et al*., 2000; Murphy *et al*.,2002; Fields *et al*., 2002), was increased in IRF2BP2^fl/fl^Lck-cre^+^ CD4^+^ T cells within all effector profiles (**Figure 5F**), explaining, at least in part, the increased expression of IFN-γ by the IRF2BP2^fl/fl^Lck-cre^+^ CD4^+^ T cells.

### In response to OVA-antigenic challenge, T cells overexpressing IRF2BP2 proliferate less than the control T cells, while Tregs are in higher frequencies

To evaluate the secondary immune response towards a specific antigen in transgenic mice, IRF2BP2^fl/fl^Lck-cre^+^ and IRF2BP2^fl/fl^Lck-cre^-^ mice were *in vitro* sensitized with ovalbumin plus Freund complete adjuvant (**Figure 6A**). After 10 days, the draining lymph nodes were taken, and the T cells were either directly analyzed (**Figure 6B-F**) or were re-stimulated *in vitro* with OVA for 72 hours (**Figure 6G-H**).

**Figure 6.**
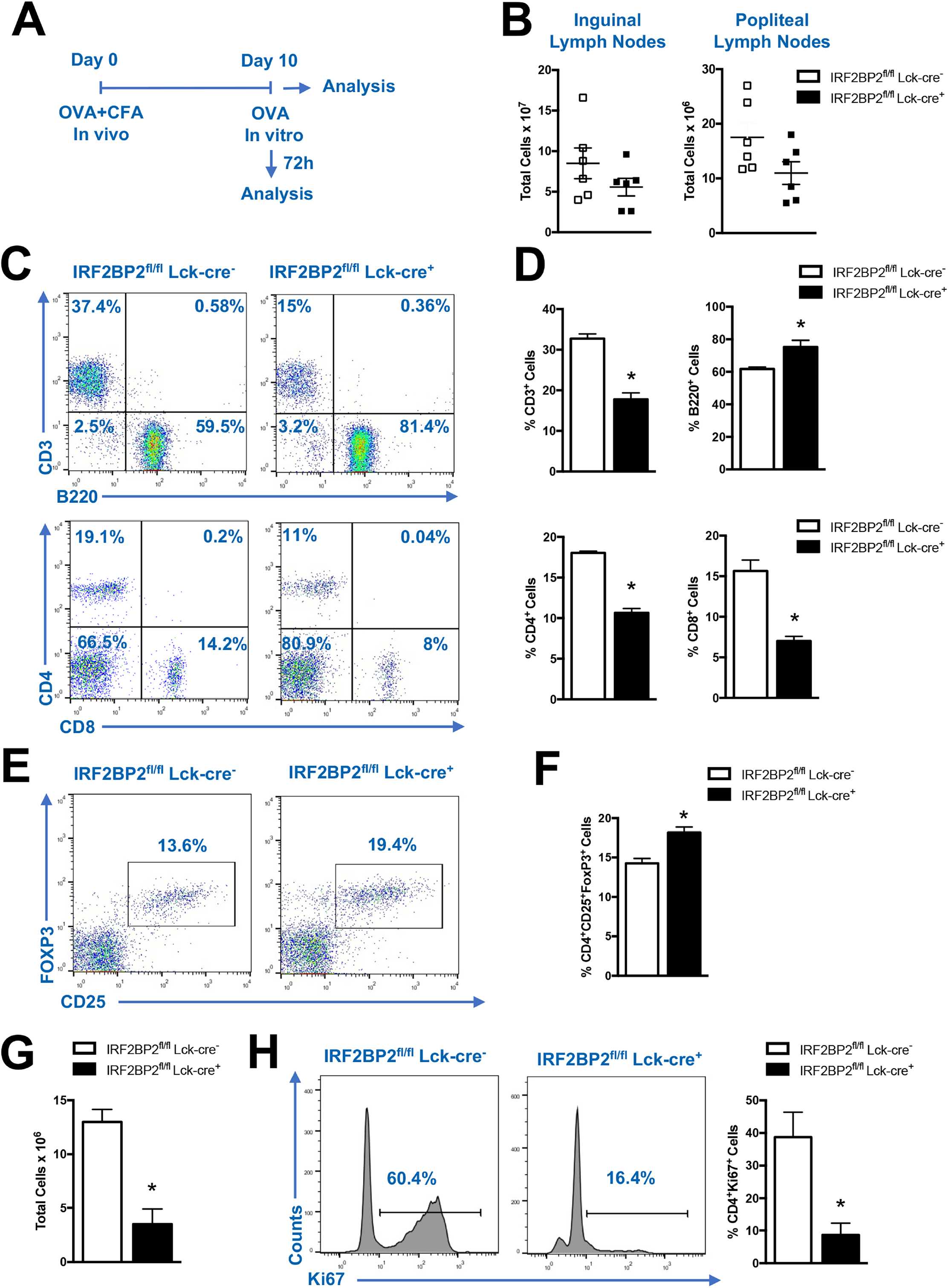
Sub-populations of lymphocytes from OVA-sensitized IRF2BP2^fl/fl^Lck-cre^+^ and IRF2BP2^fl/fl^Lck-cre^-^ mice. **A**. Schematic representation of the experiment workflow. **B**. 10 days after OVA sensitization, the mice were sacrificed and the total number of cells in the draining lymph nodes was determined by Trypan blue exclusion. **C-F**. The cells from the draining lymph nodes were stained for anti-CD3, -B220, -CD4 and -CD8 **(C-D)**; or for anti-CD4, -CD25 and - FoxP3 **(E-F). G**. The cells from the draining lymph nodes were *in vitro* stimulated with OVA (0.5 μg/mL) for 72 hours. The graph shows the total number of cells determined by Trypan blue exclusion. **H**. The cells depicted in **G** were stained for anti-CD4 and anti-Ki67. The graph on the right shows the percentage of CD4^+^Ki67^+^ cells. All the graphs show average ± SD. (*) p<0.05. Representative of 2 independent experiments (n=5).

The total number of cells obtained from the draining lymph nodes of IRF2BP2^fl/fl^Lck-cre^+^ mice was smaller than the number from control mice (**Figure 6B**). In accordance with the data obtained with naïve mice, the percentage of T cells from mice challenged with OVA overexpressing IRF2BP2 were smaller than the percentage of cells from the control mice (**Figure 6C-D**), while the Tregs were at higher frequency (**Figure 6E-F**). We also assessed CD4^+^ T cell proliferation after 72 hours of re-stimulation with OVA and we observed a decrease in total number of cells (**Figure 6G**) and a remarkable reduction in IRF2BP2^fl/fl^ Lck-cre^+^ CD4^+^ T cell proliferation when compared to the control CD4^+^ T cells (16.4% versus 60.4%, respectively - **Figure 6H**), confirming our previous results that IRF2BP2 represses T cell proliferation (Secca *et al*., 2016). Moreover, the increase in Treg percentage observed in the OVA-sensitized IRF2BP2^fl/fl^Lck-cre^+^ mice might also account for the decrease in T cell proliferation observed here and explain the smaller number of total T cells in comparison to control mice.

### Mice overexpressing IRF2BP2 in T cells exhibit reduction of clinical and inflammatory signals of EAE

So far, we showed that the overexpression of IRF2PB2 decreases T cell proliferation (**Figure 6H** and Secca *et al*., 2016), and increases IFN-γ production (**Figure 5**). We then attempted to evaluate whether the overexpression of IRF2BP2 has a role during the EAE development (**Figure 7A** and *Methods*), starting by analyzing the clinical score of the disease. Mice overexpressing IRF2BP2 presented mild signs of EAE, showing lower clinical scores compared to control mice throughout the entire course of the experiment (**Figure 7B**). The spinal cord of mice overexpressing IRF2BP2 sacrificed at the peak of the disease (day 18 after EAE induction) displayed less inflammatory cell infiltrate judged either by histology (**Figure 7C**) or by counting total lymphocytes (**Figure 7D**); and the number of Tregs (CD4^+^CD25^+^FoxP3^+^) found in the spleen was higher in mice overexpressing IRF2BP2 (**Figure 7E**), whereas the total number of CD4^+^ T cells didn’t change (*not shown*), indicating an inverse association between the frequency of Tregs and the degree of inflammation during EAE development.

**Figure 7.**
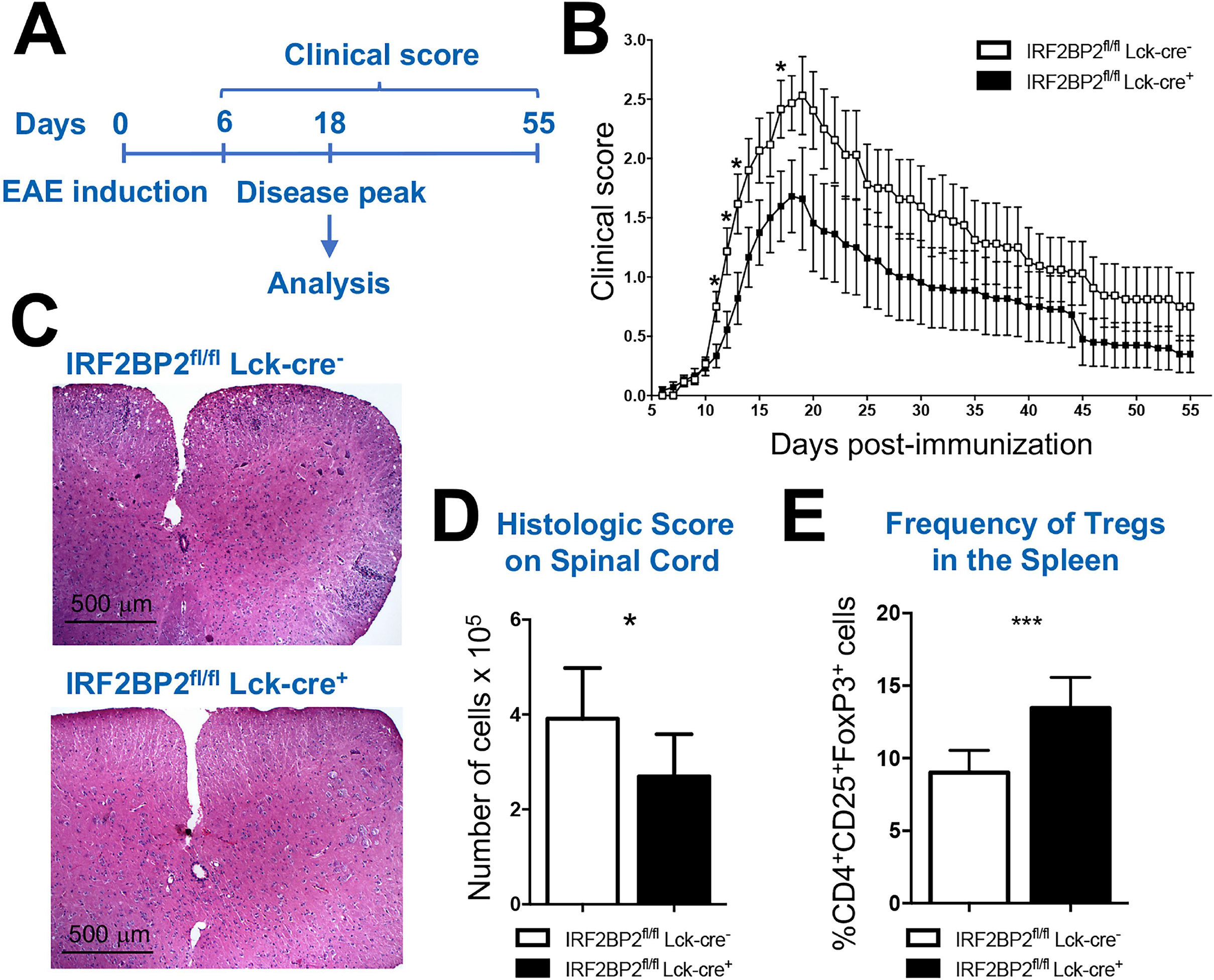
Induction of EAE in IRF2BP2^fl/fl^Lck-cre^-^ and in IRF2BP2^fl/fl^Lck-cre^+^ mice. **A**. Schematic representation of the experiment. Mice were immunized subcutaneously on the tail with MOG peptide (100 μg), complete Freund adjuvant (CFA), Mycobacterium tuberculosis (4 mg/mL) and an intraperitoneal injection with *B. pertussis* toxin (300 ng). Another round of *B. pertussis* was administrated 48 h after immunization. **B**. Disease progression and scores. The mice were monitored daily and scored clinically as described in methods. N=15 for IRF2BP2^fl/fl^Lck-cre^-^ and 18 for IRF2BP2^fl/fl^Lck-cre^+^. (*) p<0,05. **C**. Histology of the spinal cord obtained from acute EAE. Representative images of the groups 18 days after EAE induction. Transversal sections from spinal cord stained with H&E. Objective: 40X. Inflammatory cell infiltrates are in blue. **D**. Histological analysis considering the inflammatory cell infiltrate on spinal cord. For cell quantification, fields of the histological sections were analyzed using the ImageJ software and the number of cells estimated for every 50,000mm. (*) p<0,05. **E**. Frequency of Treg (CD4^+^CD25^+^FoxP3^+^) cells in the spleen of IRF2BP2^fl/fl^Lck-cre^-^ or IRF2BP2^fl/fl^Lck-cre^+^ mice after EAE induction. (***) p< 0,001.

## Discussion

The long-term aim of our laboratory is to better understand the molecular mechanisms involved in T cell activation and proliferation. Our interest in IRF2BP2 function began in our group with a yeast-two hybrid experiment to identify new partners of the transcription factor NFAT, where more than 50% of the identified clones were IRF2BP2 (Carneiro *et al*., 2011). Our subsequent studies showed that overexpression of IRF2BP2 inhibits T cell activation (Carneiro *et al*., 2011; Secca *et al*., 2016) and proliferation (Secca *et al*., 2016). The transgenic knock-in mice overexpressing IRF2BP2 in T lymphocytes constructed for this study (**Figure 1**) confirmed those results, showing a decrease in the T cell compartment when compared to control mice (**Figure 1D, 2C-F** and **6B**) and a clear reduction in T cell proliferation after secondary stimulation with OVA (**Figure 6G**).

Regarding the mechanisms by which IRF2BP2 represses T cell proliferation, while our previous results demonstrated an intrinsic role for IRF2BP2, the IRF2BP2^fl/fl^Lck-cre^+^ mice extended our knowledge, showing that IRF2BP2 is also involved in Treg function, which in turn, can repress the proliferation of surrounding cells. Tregs represent a specialized lineage of T cells essential for the immune system homeostasis (Tang & Bluestone, 2008), whose main function is the suppression of T cell responses to self-antigens (Josefowicz *et al*., 2012, Sakaguchi *et al*., 2008), preventing autoimmune diseases (Fontenot *et al*., 2003, Khattri *et al*., 2003). CD4^+^ T cells from IRF2BP2^fl/fl^Lck-cre^+^ mice upregulated the expression of the Treg specific transcription factor *Foxp3* (**Figure 4**) and exhibited a higher percentage of Tregs, either in naïve conditions (**Figure 3**), after OVA-sensitization (**Figure 6E-F**) or after EAE induction (**Figure 7E**). Therefore, this increase in Treg frequency might be directly contributing to the inhibition of cell proliferation and explaining the reduced number of T cells observed in this transgenic mouse. Clinically, IPEX (Immune dysregulation, polyendocrinopathy, enteropathy, X-linked) syndrome, caused by mutations in FoxP3, is a prototype of disorders affecting the regulation of immune responses (Bennet et al, 2001). Baxter et al (2021) have shown that 39% of IPEX-like patients - meaning patients with similar symptoms but without mutations in FoxP3 - carry mutations in 1 of 27 immuno-regulatory genes, being *irf2bp2* among them (Baxter et. al, 2021), supporting a role for IRF2BP2 in IPEX-like disease.

Another important gene upregulated in IRF2BP2^fl/fl^Lck-cre^+^_CD4^+^ T cells in our RNA-seq experiments was IFN-γ (**Figure 4**), a cytokine pivotal for Th1 differentiation and Th2 inhibition (Fields *et al*., 2002, Murphy *et al*., 2002, Zhu *et al*., 2009). Surprisingly, production of IFN-γ was increased in differentiated IRF2BP2^fl/fl^Lck-cre^+^ T cells, regardless of whether the effector profile was Th1, Th2, Th17 or the default differentiation (**Figure 5**). As shown by control cells, the production of specific cytokines is tightly regulated and exclusive to each profile after differentiation (**Figure 5**, Murphy *et al*., 2002), which does not occur with IFN-γ production of IRF2BP2^fl/fl^Lck-cre^+^ CD4^+^ T cells (**Figure 5**).

It is well described that recently activated cells begin to produce both Th1 and Th2 cytokines at low levels, a state called Th0. This state persists until a critical threshold of extrinsic promoting signals, such as cytokines, reach a level when the cell quickly shifts into a high level steady state of the transcription factors Tbet, GATA-3 or RORγt, which will determine a Th1, Th2 (Murphy *et al*., 2002) or Th17 differentiation (Mangan *et al*., 2006). These specific transcription factors are involved in chromatin changes at specific cytokine loci, such as histone acetylation of IFN-γ locus by Tbet during Th1 differentiation (Fields *et al*., 2002). The cell then remains producing high levels of one of those transcription factors with no expression of the others even if the initial Th signal is reduced below the original threshold level. Only when the Th signal is reduced to a level below the original activation threshold the cell falls back to the Th0 state (Murphy *et al*., 2002, Fields *et al*., 2002). The increase in IFN-γ production by the IRF2BP2^fl/fl^Lck-cre^+^ CD4^+^ T cells differentiated to either Th2 or Th17 differentiation clearly indicates a Th0 profile, which is accompanied by an increased expression of the transcription factor Tbet (**Figure 5F**), indicating a dysfunction during the closing of the IFN-γ locus. Why Tbet is deregulated and whether other epigenetic mechanisms, such as demethylation of the IFN-γ promoter are involved, remains to be elucidated. Notably, CIITA, a known recruiter of histone acetyltransferases induced by IFN-γ (Beresford & Boss, 2001), is upregulated in our IRF2BP2-transgenic CD4^+^ T cells (**Figure 4C**, *Transcription Factors*).

IFN-γ is also an important Th1 cytokine typically involved in EAE development (Van Den Hoogen, 2017; Olsson, 1995). It upregulates MHC class II expression that facilitates migration of T cells into the central nervous system (CNS) (Horwitz *et al*,, 1997), playing an important role in the activation of encephalitogenic T cells and CNS-inflammation. IFN-γ-producing Th1 cells enter the CNS and promote the establishment of an inflammatory microenvironment, further facilitating the recruitment of other effector cells, such as Th17, which likewise can produce IFN-γ (Sonar *et al*., 2017). Nevertheless, blocking IFN-γ exacerbates EAE because IFN-γ is necessary to convert conventional CD4^+^CD25^-^ T cells into Tregs (Wang *et al*., 2006). IFN-γ also recruits macrophages and Tregs across the blood-CSF barrier at the inflamed spinal cord (Kunis *et al*., 2013), helping in the repair of the injured spinal cord, meaning that IFN-γ has both inflammatory and regulatory roles in a stage-specific manner in multiple sclerosis and EAE (Arellano *et al*., 2015; Sonar *et al*., 2017). Interestingly, some genes induced by IFN-γ known for playing a role in EAE were upregulated in IRF2BP2^fl/fl^Lck-cre^+^ T cell-RNA-seq, such as the chemokines CCL4 and CCL5 (**Figure 4C**, *Cytokines*, Sonar *et al*., 2017).

The signs of EAE were milder in mice overexpressing IRF2BP2 versus control mice, with less inflammatory T CD4^+^ cell infiltrate in the spinal cord, and increased number of Tregs, suggesting that the reduction in the symptoms of EAE is likely due to an increased immunoregulatory activity. Besides the roles described before, another pivotal function of Tregs is to limit immune responses to infections, decreasing the corresponding tissue damage (Josefowicz *et al*., 2012, Arpaia *et al*., 2015), and we believe that this is the mechanism by which IRF2BP2 overexpression decreases EAE symptoms (**Figure 7**). It was reported that Tregs can produce IFN-γ under Th1-polarization conditions (Oldenhove *et al*., 2009; Zhao *et al*., 2012) and that these cells represent the first line of Tregs that suppress an initial immune response (Daniel *et al*., 2014).Arruda *et al*., 2015, showed that besides IRF2BP2 and FoxP3 being down-regulated in T cells from patients with multiple sclerosis, IRF2BP2 levels are restored in patients in clinical remission, indicating that IRF2BP2 controls T cell-mediated disease inflammation (Arruda *et al*., 2015), supporting our hypothesis that IRF2BP2 plays a role as an immunoregulatory protein. Further experiments will be needed to establish whether the number of Tregs is also increased in the spinal cord after EAE-induction, and whether the production of IFN-γ is increased in the infiltrating lymphocytes.

A regulatory/anti-inflammatory role for IRF2BP2 has been demonstrated before, inhibiting the pro-inflammatory M1 and promoting the anti-inflammatory M2 phenotype in macrophages (Chen *et al*., 2015). Likewise, the loss of IRF2BP2 in microglia increases the production of inflammatory cytokines after ischemic brain injury (Cruz *et al*., 2017). IRF2BP2 also increases PD-L1 expression (Bhadra *et al*., 2011; Soliman *et al*., 2014) and the mechanism seems to involve IFN-γ (Wu *et al*., 2018), which increases the association between IRF2BP2 and IRF-2, releasing IRF-2 from the PD-L1 promoter, increasing its expression. Interestingly, IFN-γ also induces PD-1 expression (Cheng *et al*., 2007) and *Pdcd1* (which codifies PD-1) is increased in IRF2BP2^fl/fl^Lck-cre^+^ T cells (**Figure 4C**, *Plasma Receptors*). Collectively, these findings provide indications that IFN-γ works alongside with IRF2BP2 to restrain the immune response, which might have implications in the establishment of autoimmune diseases and cancer.

## Materials and Methods

### Construction of a conditional IRF2BP2 *knock-in* mouse

The plasmid pLIRES-EGFP-mIRF2BP2 (Secca *et al*., 2016) was used to amplify the murine IRF2BP2 cDNA. The AscI restriction site, a kozak sequence and a C-terminal myc-tag were included in the primers. The primers used were: pr F:5’ GGC GCG CCa aAC Cat ggc cgc ggc ggt ggc and pr R: 5’ GGC GCG CCt caC AGA TCC TCT TCT GAG ATG AGT TTT TGT TCc gag tct ctc tcc ttc ttc. The PCR was performed using Phusion DNA polymerase (NEB) and DMSO 10%. The PCR product was cloned into TOPO vector (Invitrogen) after Poly-A-tailing reaction and further released using the AscI restrinction sites. The insert was sequenced and cloned in the AscI site of CTV-vector (kindly provided by Xiao Changchun – The Scripps Research Institute, CA, US), also available at addgene (plasmid 15912) (http://www.addgene.org/15912/) using the Roche Rapid ligation kit (Roche). The correct insert orientation was confirmed by NheI and SmadI (NEB) restriction pattern. The final vector, CTV-mIRF2BP2-myc, was linearized with SgfI (Promega) and electroporated into the embryonic stem (ES) cell line PRX. After 3 weeks, the gDNA of the G418-resistant ES colonies was extracted to confirm the transgene integration. The ES cell clones were first screened out by PCR (pr F: 5’ gtg gtt tgt cca aac tca tca and pr R 5’ tcc gtg aag tcc cag atc at), and confirmed by Southern blot. Confirmed positive ES cell clones were injected into albino C57BL/6 blastocysts to generate chimerical mice, which were then bred with C57BL/6 mice for germline transmission. ES cell culture and microinjection were performed by the Mouse Genetics Core at The Scripps Research Institute in La Jolla, CA, US. IRF2BP2-floxed-targeted germline transmitted mice (IRF2BP2^fl/fl^) bred with Lck-Cre mouse (Taconic’s lab), in which the Cre recombinase is expressed under the control of Lck promoter, to induce germline overexpression of IRF2BP2 in T cells, to generate the final mouse: IRF2BP2^fl/fl^Lck-cre^+^. All mice were maintained in specific-pathogen-free barrier facilities and used according to protocols approved by the La Jolla Institute for Allergy and Immunology animal care and use committees. The final mice were transferred to the Brazilian National Cancer Institute (INCA, RJ, Brazil) under the Brazilian Government’s ethical and animal experimental regulations. All mice used for the experiments were between 8-12 weeks old, and all the experiments were approved according to the animal welfare guideless of the Ethics Committee of Animal Experimentation of INCA.

### Southern blot

The gDNA from Th1 cells from C57BL/6 mice was used as template to amplify a 490 bp probe, inside the ROSA26 genomic loci, using the primers: pr F: 5’ TTC ATG TCA CTT GCA CTG GGA A and pr R 5’ AGT GAG TAT ACA AGG CGG CAG. The probe was labeled with the Takara Ladderman kit (cat. 6046) using the manufacture’s recommendations. 10 μg of genomic DNA from ES clones was digested overnight with EcoRI. The following day, the digested gDNA was run in an 0.8% agarose gel for 6h at 70V. The gel was washed with distilled water for 5 minutes, crosslinked with 700 mJ UV radiation, washed for 10 minutes with Southern Transfer Buffer (0.4M NaOH, 1M NaCl) and transferred overnight to a Hybond membrane pre-wet with Southern Transfer Buffer. On the next day, the membrane was neutralized in 0.5 M Tris pH 7 for 5 minutes, was dried overnight and then crosslinked with 1200 mJ of UV radiation. The membrane was wet with 2X SSC (0.3M NaCl, 30 mM Trisodium citrate, pH 7) and transferred to a hybridization tube, followed by incubation with agitation with 2X SSC for 10 minutes in an oven at 65C. The SSC buffer was discarded and exchanged by 12mL of Hybridization buffer, 1h 65C. The buffer was discarded and replaced by 12mL of Hybridization buffer in the presence of the labeled probe (10^6^ CPM/mL) overnight at 65C. The blot was washed twice with 100 mL 2X SSC for 10 minutes at 65C and twice with 100mL 2X SSC 0.1% SDS. The blot was exposed for 2 hours at -80C using a KODAK BioMax^®^ Maximum Sensitivity (MS) Autoradiography Film.

### Quantitative Real-time RT-PCR

Total RNA was extracted from 500,000 CD4^+^ T cells from IRF2BP2^fl/fl^ Lck-cre^+^ or IRF2BP2^fl/fl^ Lck-cre^-^ mice using Qiagen RNeasy mini kit. The RNA from activated or differentiated T cells was extracted using TRIzol (Thermo Fisher Scientific). cDNA was synthesized using Superscript reverse transcriptase and oligo(dT) primers (Invitrogen), and gene expression was examined with Step One Plus (Applied Biosystems) using TaqMan^®^ Real Time PCR Assay. The TaqMan probes used were Mm01239804_g1 (IRF2BP2), Mm 00450960_m1 (Tbx21) and Mm00446968_m1 (HPRT). IRF2BP2 and Tbet gene expression were normalized by the HPRT gene expression.

### Isolation and culture of T cells

Spleen, thymus and lymph nodes from IRF2BP2^fl/fl^Lck-cre^+^ or IRF2BP2^fl/fl^Lck-cre^-^ mice were harvested from 8 to 12-week-old mice. Splenic cells were first macerated and incubated with ACK solution (Gibco) for 1 min to remove the red cells and then washed 2X with 1X PBS. Naïve CD4^+^ (CD25^lo^, CD44^lo^, CD62L^hi^) T cells were purified (>95% purity) by negative selection using the Dynal Mouse CD4 Negative Isolation kit (Invitrogen). When indicated, the cells were cultured in Dulbecco’s modified Eagle’s medium (DMEM) supplemented with 10% heat-inactivated fetal bovine serum, 2 mM L-glutamine, penicillin-streptomycin, non-essential amino acids, sodium pyruvate, vitamins, 10 mM HEPES, and 50 μM 2-mercaptoethanol. Cells were plated at 10^6^ cells/ml in 6-well plates plate-coated with anti-CD3 (clone 2C11) and anti-CD28 (clone 37.51) (1 μg/ml each, BD Biosciences) by pre-coating the wells with 100 μg/ml goat anti-hamster IgG (Cappel).

### Surface Staining, Intracellular Staining and Flow Cytometry

For surface staining, the indicated cells were labelled for 15 min at RT in the dark with the following anti-mouse antibodies: anti-CD3-APC, anti-CD4 PE, anti-CD44 APC or BV421, anti-CD25 PerCP-Cy5.5, anti-CD8-PerCP-Cy5.5, anti-CD25-APC, anti-CD62L-PerCP-Cy5.5, anti-CD69-APC, anti-CD122-PE, anti-B220-PE (all BD Biosciences) and anti-CD62L APC-Cy7 (Biolegend). For cytokine and FoxP3 intracellular staining, the cells were first labeled with the indicated surface staining antibodies, then fixed and permeabilized using the Foxp3 staining buffer kit from eBioscience following the manufacture’s recommendations. For cytokine analysis, the cells were stimulated with 10 nM PMA and 1µM ionomycin for 6 hours at 37C. In the final 2 hours of stimulation, 10 µg/mL Brefeldin A (eBioscience) was added to the cell cultures. The intracellular antibodies used were: anti-FOXP3 APC and PerCP-Cy5.5, anti-IL4-APC, anti-IL-17 APC, anti-IFN-γ-PE (all BD Biosciences), and anti-Ki67-PE (eBioscience). Flow cytometry analysis was performed in a FACScan, FACSCanto, or BD FACSARIA III (Becton Dickinson) and data analyzed using the FlowJo software (Tree Star Inc).

### RNA-seq

Naive CD4^+^ T cells (CD25^lo^, CD44^lo^, CD62L^hi^) from IRF2BP2^fl/fl^Lck-cre^+^ or IRF2BP2^fl/fl^Lck-cre^-^ mice were sorted by fluorescence-activated cell sorting (FACS, BD FACS ARIA III) and left unstimulated (data not shown) or were stimulated for 1 hour with PMA (10 nM) plus ionomycin (1 μM). Total RNA was extracted from 500,000 CD4^+^ T cells using the RNeasy kit. RNA quality was evaluated with Bioanalyzer RNA pico kit (Agilent Technologies Inc, Santa Clara, CA). Poly(A)-selected RNA was amplified using the SMARTseq2 protocol (Clontech – Picelli *et al*., Nat Prot, 2014). Briefly, purified RNA was reverse-transcribed using Superscript II, Oligo dT30 VN primers and template switching primers. A pre-amplification step of 9 PCR cycles was performed using the Kapa HiFI Hoststart kit (Kapa Biosystems). The PCR product was purified using AmpureXP beads (Beckman Coulter, Danvers, MA) and 1 ng was further used for library preparation using the Nextera XT LibraryPrep kit (Illumina). Tagmented DNA was amplified with a 12 cycle PCR reaction and again purified with AmpureXP beads. Library size distribution and yield were evaluated using the Bioanalyzer high-sensitivity DNA kit. Libraries were pooled at equimolar ratio and sequenced with the rapid run protocol on the Illumina HiSeq 2500 with 50 paired end cycles.

### RNA-seq Analysis

Trimmed reads using the software Trimmomatic. (Bolger *et al*., 2014) with default options were mapped to the mouse genome (GRCm38) using STAR (Dobin *et al*., 2013) with default parameters for paired-end unstranded RNA-seq data as per developer’s manual. Htseq count (Anders *et al*., 2015) was used to assign uniquely mapped reads to genes according to the annotation in the Mus_musculus.GRCm38.83.gtf (excluding pseudogenes). Groups comparison: read counts were analyzed using the R package DESeq2 (Love *et al*., 2014) and libraries were normalized using the estimateSizeFactors function of the package. Differentially expressed genes were required to pass a log fold change threshold of 0.4 and a FDR 0.05. Heatmaps were constructed using the package pheatmap using a regularized Log2-transformed counts-per-million, z-scaled across samples.

### Imunofluorescence

Purified CD4^+^ T cells from IRF2BP2^fl/fl^Lck-cre^-^ and IRF2BP2^fl/fl^Lck-cre^+^ mice were *in vitro* activated with PMA (20 nM) and Ionomycin (2 µM) for 24 hours. After stimulation, cells were fixed with 4% paraformaldehyde (PFA) for 15 minutes at room temperature. The cells were then washed with wash buffer (1X PBS, 5% FBS and 0,5% NP40) for 5 times (5 minutes each) using slight agitation. The cells were incubated with anti-IRF2BP2 (Proteintech) for 2 hours at room temperature and then washed with wash buffer for 5 times (5 minutes each), and subsequently incubated with rhodamine-conjugated secondary antibody (KPL), followed by 5 washes with 1X PBS (5 minutes each). The cells were also stained with DAPI (10 mM) (Invitrogen) for 5 minutes. Coverslips were mounted on glass slides using Vectashield antifade mounting medium and photographed with Confocal Laser Scanning Microscope FV10i-O Olympus.

### *In vitro* cell differentiation

Purified CD4^+^ T lymphocytes were *in vitro* activated with plate-coated anti-CD3 and anti-CD28 antibodies (1µg/mL, both BD Biosciences) in the presence of specific recombinant cytokines and neutralizing antibodies: Th1 cell cultures received 5 ng/mL of recombinant IFN-γ, 5 ng/mL IL-12 and 20 µg/mL anti-IL-4 (11B11, hybridoma) Th2 cell cultures received 50 ng/mL IL-4 and 100 µg/mL anti-IFN-γ antibody (XMG.1.2, hybridoma). Th17 cell cultures received 5 ng/mL TGF-β and 50 ng/mL IL-6 recombinant and 20 µg/mL anti-IL-4 and 100 µg anti-IFN-γ (PeproTech). After 48 h, cells were removed from the TCR signal and re-cultured by doubling the area and volume in media supplemented with 20 U/ml IL-2 (Peprotech), which was added every 3 days, until day 6. Expression of cell surface receptors and cytokine production were assessed by flow cytometry.

### Gene expression analysis

Total RNA was extracted from total CD4^+^ T cells *in vitro* activated and differentiated using TRIzol (Invitrogen), followed by cDNA synthesis by RT-PCR, using the enzyme reverse transcription reaction using the Superscript II – Reverse trasncriptase (Invitrogen). Gene expression analyzes performed by real-time PCR, in the TaqMan Gene Expression Assay system (Life Technologies). We use 10 ng of cDNA, synthesized for each reaction. Probes were used specific for Tbx21 gene and Hprt (Hypoxanthine-guanine phosphoribosyltransferase). Hprt was used as the housekeeping gene for mass normalization, (Taqman Mm00450960_m1 and HPRT Mm01545399_m1). The probes used for the genes tested were conjugated to FAM fluorophores. The analysis was relative quantification of 2^-ΔΔCt^ for the levels of mRNA obtained from activated and differentiated in Th2 profile of animals IRF2BP2^fl/fl^Lck-cre^-^ in comparison with activated CD4 T cells of animals IRF2BP2^fl/fl^Lck-cre^+^. All procedures were performed according to the manufacturer’s suggested protocols.

### Sensitized animals with Ovalbumin

IRF2BP2^fl/fl^Lck-cre^-^ and IRF2BP2^fl/fl^Lck-cre^+^ mice were immunized subcutaneously in the hindfoot with a single dose of 100 μL containing 200 μg Ovalbumin (OVA, Sigma) in the presence of complete Freund’s adjuvant. After 10 days, the animals were sacrificed, the draining lymph nodes were removed and the lymphocytes were analyzed by Flow Cytometry, as previously described.

### Proliferation assay

Draining lymph nodes from IRF2BP2^fl/fl^Lck-cre^-^ and IRF2BP2^fl/fl^Lck-cre^+^ animals sensitized with OVA were harvested. The cell suspension was *in vitro* re-stimulated with OVA (0.5 μg/mL) for 72 hours in a 96-well plate (Biofil) at 37C, 5% CO_2_. The total cell number was quantified by the manual counting method with exclusion of non-viable cells with 0.4% trypan blue in saline solution. The cells were surface stained, fixed and permeabilized using the Foxp3 staining buffer kit (eBioscience) following manufacture’s recommendations, labeled with anti-Ki67 (PE) for 30 min at RT in the dark and analyzed by flow cytometry. The cells were photographed in Axionvision Observer Z1 fluorescence microscope (Zeiss, Germany).

### EAE induction

IRF2BP2^fl/fl^Lck-cre^+^ or IRF2BP2^fl/fl^Lck-cre^-^ mice at 8-10 weeks old were immunized subcutaneously (s.c.) at the base of the tail with 100 μg of myelin oligodendrocyte glycoprotein peptide (MOG35-55, Proteimax, Brazil) (MEVGWYRSPFSRVVHLYRNGK) in CFA (complete Freund’s adjuvant) containing 4 mg/mL of Mycobacterium tuberculosis H37RA (Difco, USA). Pertussis toxin (300 ng) was injected intraperitoneally (i.p.) at the day of immunization and 48 h later. Clinical evaluation of EAE was performed according to the following criteria: 0 no motor change; 0.5 partial tail paralysis; 1 tail paralysis or change in walking pattern; 1.5 partial tail paralysis and change in walking pattern; 2 tail paralysis and change in walking pattern; 2.5 partial limb paralysis; 3 paralysis of one limb; 3.5 paralysis of one limb and partial paralysis of another one; 4 complete hind-limb paralysis; 4.5 complete hind-limb paralysis and partial paralysis of the front limb; 5 animal with virtually no movement. Animals were observed for 55 days after immunization to evaluate disease development or until day 18 and were sacrificed for removal of spleens and spinal cords.

### Histopathology of EAE

Spinal cords of the mice were fixed in a solution containing 20% dimethylsulfoxide and 80% methanol for 6 days at -80°C, then for 24 hours at -20°C. Samples were then packed in Paraplast^®^, cut into pieces of 5 μm thickness and stained with hematoxylin and eosin. Sections were evaluated for histopathological changes, such as inflammatory infiltrate and white matter lesions using an optical microscope (Olympus BX41) and a camera (Moticam 2500) for imaging. For cell quantification, we used the ImageJ software in which representative histological sections were analyzed and measured. Bar graphs were plotted by Prisma 6.0 software.

### Statistical analysis

Statistical analysis used unpaired Student’s *t* test for the single comparison using the program GraphPad Prism (GraphPad software, San Diego, CA, USA). *P* ≤ 0.05 were considered statistically significant.

## Author contributions

G.P.M., H.B., A.R., A.M.C.F and J.P.B.V. designed research; G.P.M., B.O-V., N.P.-R., B.P., and C.S. performed research; M.B. and E.G-A. analyzed RNA-seq data; G.P.M., B.O-V., N.P-R., B.C.P., C.S., H.B., A.M.C.F. and J.P.B.V. analyzed data; and G.P.M. wrote the paper.

## Acknowledgments

We thank the La Jolla Institute (LJI) Flow Cytometry Core Facility for help with cell sorting; the La Jolla Institute (LJI) Bioinformatics Core, and Jeremy Day of the Sequencing Core for help with RNA-seq. This work was supported by grants to JPBV from CNPq (408127/2016-3 and 307042/2017-0), FAPERJ (203.007/2016) and INCT-Cancer (573806/2008-0 and 170.026/2008). The HiSeq 2500 was funded by NIH S10OD016262 and the FACSAria Cell Sorter by the S10RR027366. G.P.M. and C.S. were supported by CNPq fellowship; B.O.V., N.P.R. and B.C.P. were supported by a CAPES fellowship.

